# CpelTdm.jl: a Julia package for targeted differential DNA methylation analysis

**DOI:** 10.1101/2020.10.17.343020

**Authors:** Jordi Abante, John Goutsias

## Abstract

**Motivation:** Identifying regions of the genome that demonstrate significant differences in DNA methylation between groups of samples is an important problem in computational epigenetics. Available methods assume that methylation occurs in a statistically independent manner at individual cytosine-phosphate-guanine (CpG) sites or perform analysis using empirically estimated joint probability distributions of methylation patterns at no more than 4 contiguous CpG sites. These approaches can lead to poor detection performance and loss of reliability and reproducibility due to reduced specificity and sensitivity in the presence of insufficient data.

**Results:** To accommodate data obtained with different bisulfite sequencing technologies, such as RRBS, ERRBS, and WGBS, and improve statistical power, we developed CpelTdm.jl, a Julia package for targeted differential analysis of DNA methylation stochasticity between groups of unmatched or matched samples. This package performs rigorous statistical analysis of methylation patterns within regions of the genome specified by the user that takes into account correlations in methylation and results in robust detection of genomic regions exhibiting statistically significant differences in methylation stochasticity. CpelTdm.jl does not only detect mean methylation differences, as it is commonly done by previous methods, but also differences in methylation entropy and, more generally, between probability distributions of methylation.

**Availability and Implementation:** This Julia package is supported for Windows, MacOS, and Linux, and can be freely downloaded from GitHub: https://github.com/jordiabante/CpelTdm.jl.

## Introduction

Differential analysis of DNA methylation using bisulfite sequencing data is an important task in epigenetic studies. Although many statistical approaches have been developed for this task (Landan *et al*., 2012; Li *et al*., 2014; Robinson *et al*., 2014), most methods do not directly account for correlations in methylation, while others cannot reliably estimate joint probability distributions (PDMs) of methylation patterns, due to the lack of sufficient data to properly estimate a geometrically growing number of parameters, leading to reduced specificity and sensitivity as well as loss of reliability and reproducibility (Jenkinson *et al*., 2017, 2018). To deal with these issues, Jenkinson *et al*. (2017) developed informME, a method for genome-wide analysis of methylation stochasticity that uses concepts from statistical physics and information theory. To achieve a computationally manageable approach to genome-wide methylation analysis, informME characterizes DNA methylation using the probabilities of methylation levels (percentage of methylated CpG sites) within non-overlapping genomic units composed of 150 bp each, and performs analysis at the resolution of one genomic unit. Consequently, this method does not allow direct comparisons between PDMs, which can provide valuable insights into mechanisms underlying the formation of methylation patterns (Landan *et al*., 2012). In addition, informME employs a hypothesis testing approach based on a null distribution that is computed from control samples, thus introducing directionality in the analysis that is not always desirable. The genome-wide nature of methylation analysis performed by informME may also reduce its power to reject the null hypothesis when the alternative hypothesis is true, potentially leading to false negatives, a problem that can be remedied by performing targeted analysis on specific genomic regions of interest.

To address these issues, we developed a differential method for methylation pattern analysis by using a recently introduced correlated potential energy landscape (CPEL) model (Abante *et al*., 2020) and by employing non-directional permutation methods for hypothesis testing. This approach, which is implemented in the freely available Julia package CpelTdm.jl, allows a user to perform targeted differential methylation (TDM) analysis of the stochastic behavior of methylation patterns between two groups of *unmatched* or *matched* bisulfite (RRBS, ERRBS, or WGBS) sequencing data in a statistically rigorous manner.

### Approach

CpelTdm.jl involves several steps; see Fig. 1 and the Methods section. First, it uses a CPEL approach to characterize DNA methylation within analysis regions that are determined from a user-specified single-track input BED file. This approach models the stochastic methylation state at all CpG sites within an analysis region via a PDM specified by a small number of parameters, which accounts for correlations among the methylation states at contiguous CpG sites. It subsequently uses independent and possibly partially observed bisulfite sequencing reads of the methylation state and estimates the values of the PDM parameters via maximum-likelihood. Finally, it uses the estimated CPEL models to perform differential analysis between two groups of unmatched or matched data samples by testing the null hypothesis that no differences in the information content of the methylation state are present in a given genomic region. This is done by defining appropriate differential test statistics in terms of the mean methylation level (MML), which evaluates the fraction of methylated CpG sites within an analysis region, the normalized methylation entropy (NME), which quantifies the amount of methylation stochasticity (pattern heterogeneity), and the coefficient of methylation divergence (CMD), which quantifies PDM differences using the information-theoretic notions of Kullback-Leibler divergence and cross entropy; see Figs. 2 & 3 and the Methods section.

**Figure 1.**
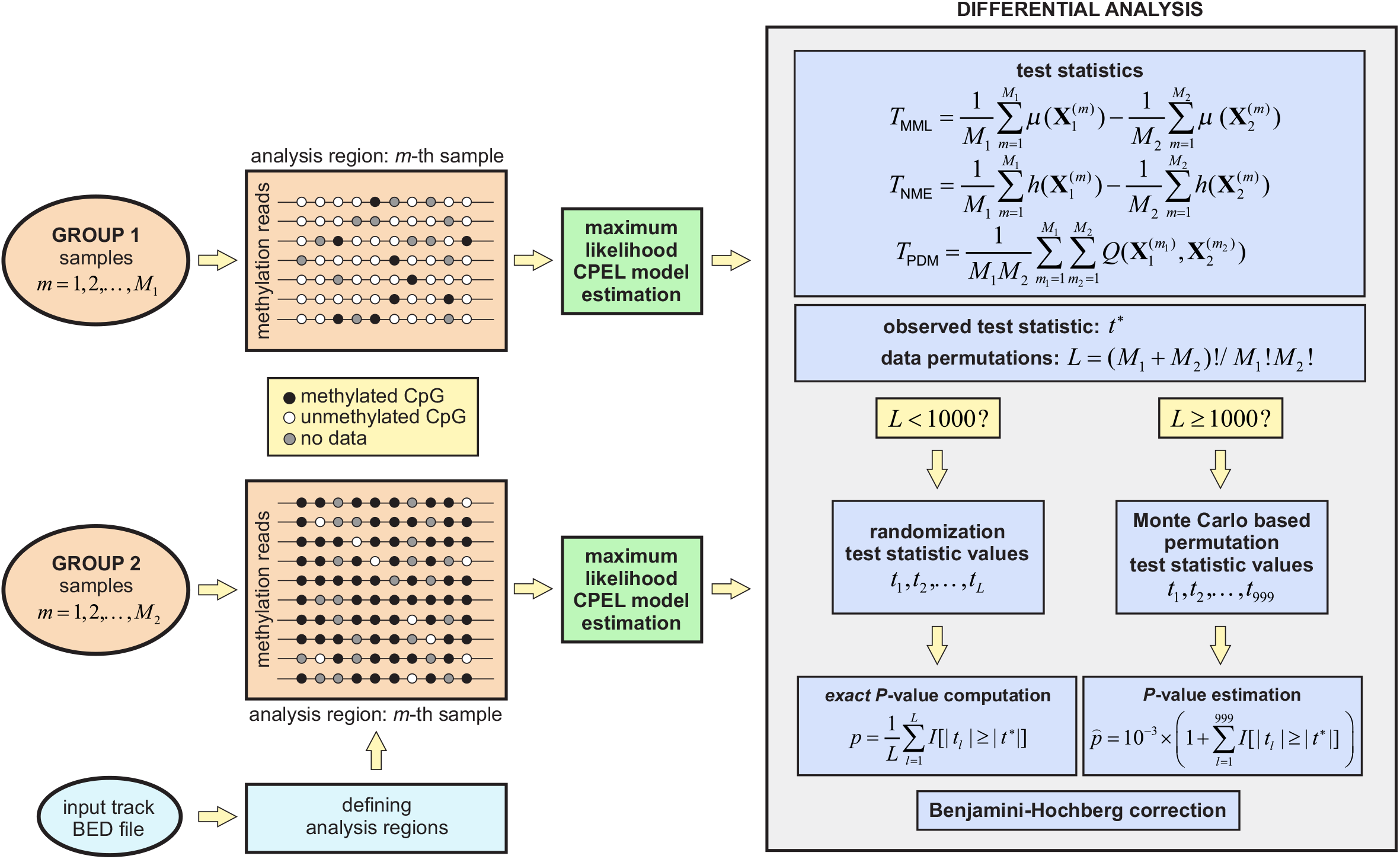
Flowchart of CpelTdm.jl for differential methylation analysis between two groups of unmatched samples in terms of the mean methylation level (MML) *μ*, the normalized methylation entropy (NME) *h*, and the coefficient of methylation divergence (CMD) *Q*. A similar approach is employed in the case of matched samples.

**Figure 2.**
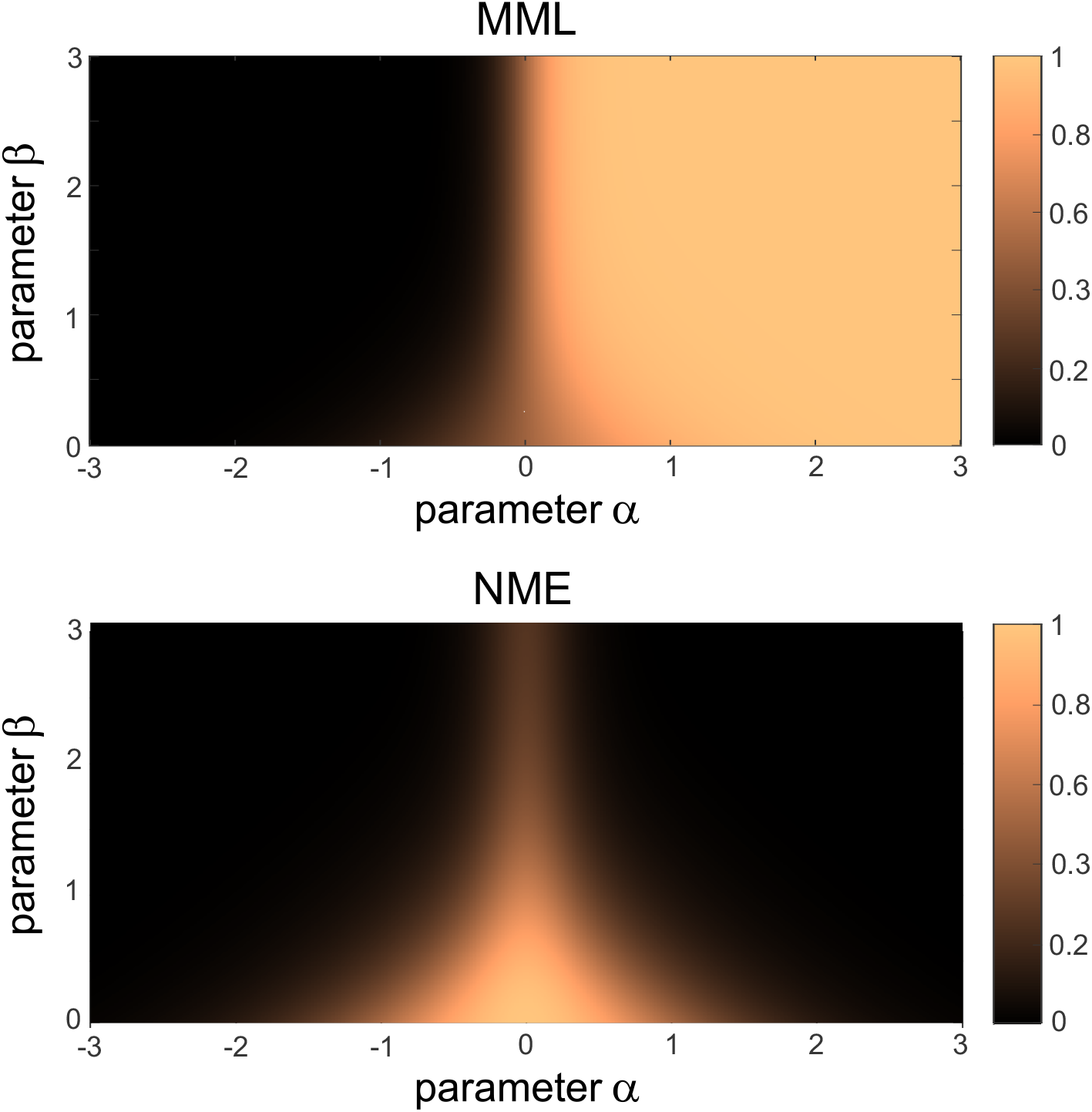
Heat maps of the mean methylation level (MML) and the normalized methylation entropy (NME) within an analysis region containing 4 CpG sites whose methylation state is characterized by a CPEL model with two parameters −3 ≤ *α* ≤ 3 and 0 ≤ *β* ≤ 3. Smaller negative values of *α* result in decreased MML values, indicating that fewer CpG sites are methylated on the average, whereas larger positive values result in increased MML values, indicating that more CpG sites are methylated on the average. This behavior is associated with increasingly reduced NME values in both cases, and is exacerbated with larger values of *β* due to increased correlation. When *α* = *β* = 0, each CpG site is methylated independently from each other with 50% probability, resulting in an equal number of CpG sites to be methylated and unmethylated on the average (MML = 0.5) and maximum methylation entropy (NME = 1).

**Figure 3.**
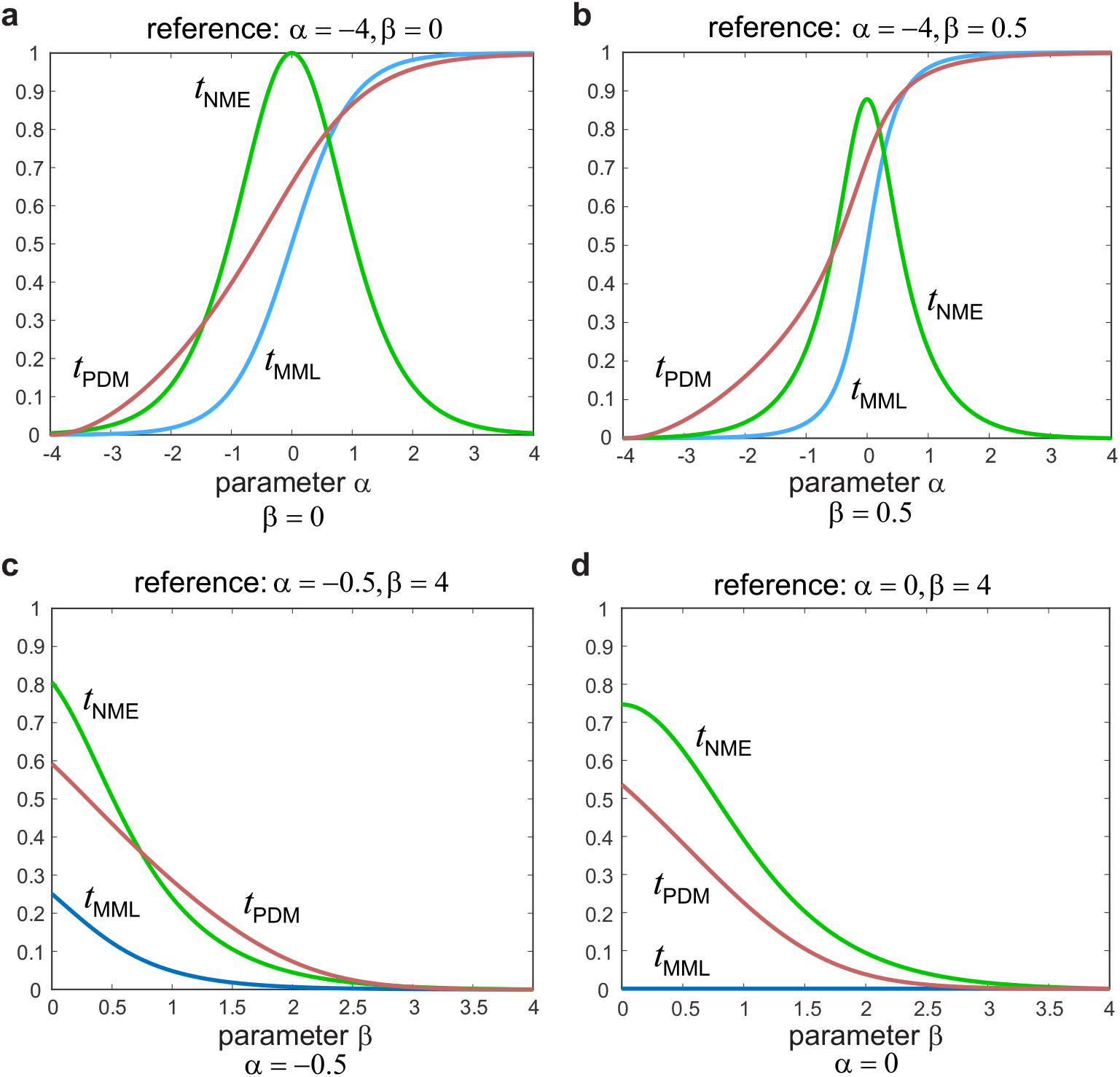
Examples of differential test statistic values *t*_MML_, *t*_NME_, and *t*_PDM_ between a reference and a test analysis region containing 4 CpG sites whose methylation state is characterized by a CPEL model with two parameters *α* and *β*. (a) Reference model: *α* = −4, *β* = 0; test models: −4 ≤ *α* ≤ 4, *β* = 0. (b) Reference model: *α* = −4, *β* = 0.5; test models: −4 ≤ *α* ≤ 4, *β* = 0.5. (c) Reference model: *α* = −0.5, *β* = 4; test models: *α* = −0.5, 0 ≤ *β* ≤ 4. (d) Reference model: *α* = 0, *β* = 4; test models: *α* = 0, 0 ≤ *β* ≤ 4. The CpG sites in the reference analysis region are unmethylated in (a), (b), and (c), whereas half of them are methylated in (d) on the average. Notably, the test region transitions from being unmethylated to being methylated as *α* increases in (a) and (b), and to being partially methylated as *β* decreases in (c). Interestingly, no MML difference is present in (d), despite differences in NME and PDM. Note also that methylation is uncorrelated in (a), and this is also true in (c) and (d) when *β* = 0. On the average, most CpG sites in the test analysis region are methylated (*t*_MML_ > 0.5) in (a) and (b) when *α* > 0 and unmethylated (*t*_MML_ < 0.5) when *α* < 0. Moreover, the differential test statistic *T*_NME_ achieves its maximum value when *α* = *β* = 0 in (a), corresponding to a CPEL model which assumes that the CpG sites in the test analysis region are independently methylated with 50% probability.

The main challenge in performing hypothesis testing is computing the null distribution of a test statistic from given data when a theoretical distribution is not available. To address this problem and for computational efficiency, CpelTdm.jl performs permutation testing using one of two approaches depending on the number *L* of all possible data permutations: (exact) randomization testing is performed when *L* < 1000, and (approximate) Monte Carlo based permutation testing is performed when *L* ≥ 1000; see the Methods section. Both approaches properly control the Type I error (false positives), in the sense that the probability of such an error is no larger than the test’s significance level. The resulting *P*-values are finally corrected for multiple hypothesis testing using the Benjamini-Hochberg procedure for FDR control.

## Results

Using simulations, we investigated the effectiveness of CpelTdm.jl for correctly identifying significant differences in methylation stochasticity between two groups containing 5 or 7 unmatched and 6 or 10 matched samples each. We considered a small genomic region with 4 CpG sites and characterized methylation stochasticity within the region by a CPEL model with parameters α and *β* of known ‘true’ values leading to the following four scenarios: no methylation differences, differences in MML and PDM, differences in NME and PDM, and differences in MML, NME, and PDM.

We generated methylation data in agreement with the number of samples associated with each group. Each data sample was associated with *R* methylation reads obtained by sampling the ‘true’ CPEL model corresponding to each group, where the value of *R* was randomly determined to be between 10 and 30 with equal probability. Using these data, we computed an ‘estimated’ CPEL model for each sample, from which we calculated test statistic and corresponding *P*-values in a specific group comparison. Finally, by repeating this process 1000 times, we derived empirical estimates of the cumulative distribution functions of the *P*-values for each differential test statistic; see Figs. 4–7. The results showed the ability of CpelTdm.jl to detect statistical significance at a high rate (at least 95% of the time) while keeping the probability of false detections below 5% in all cases, thus demonstrating its reliability for correctly performing differential methylation analysis.

**Figure 4.**
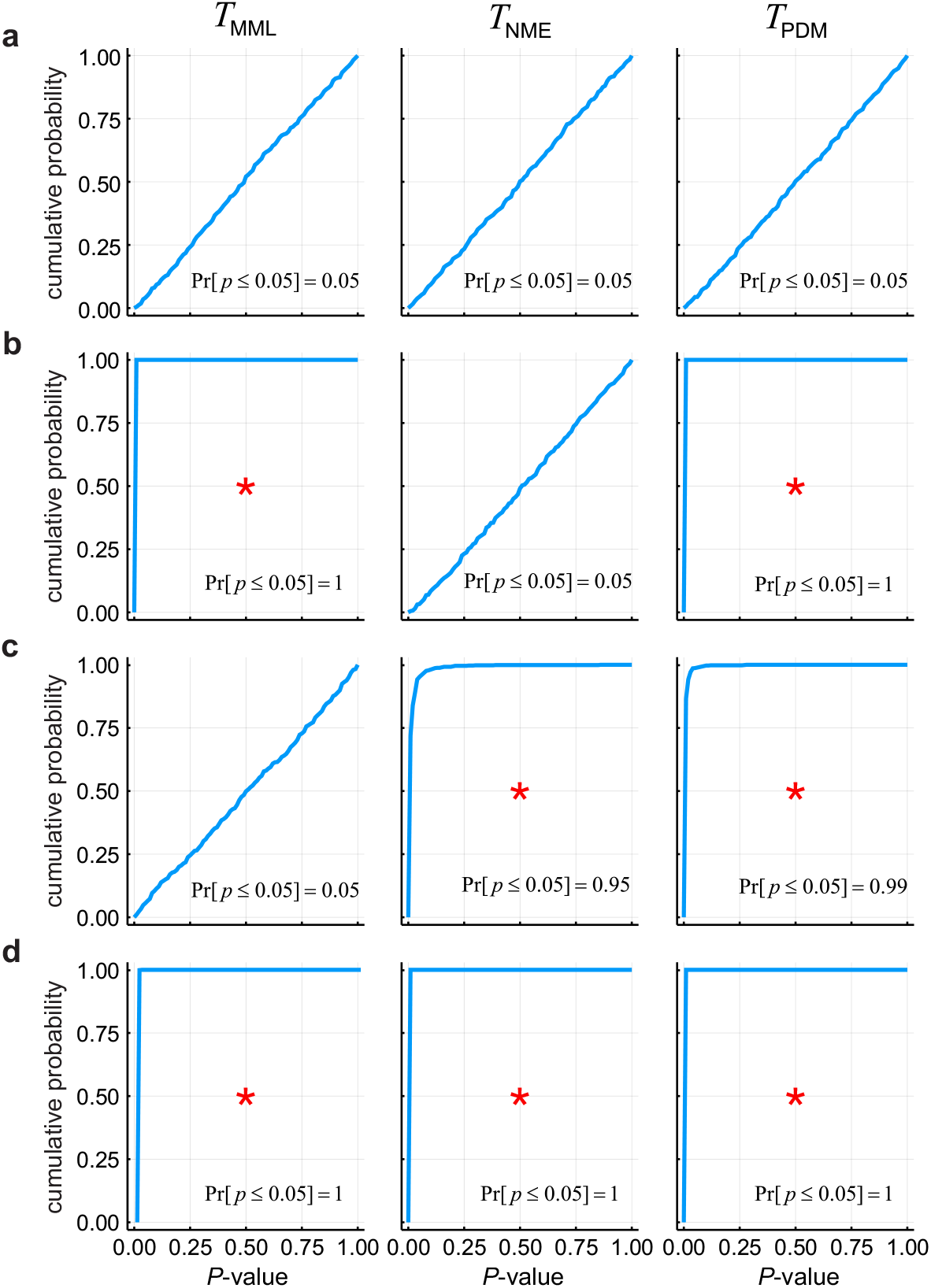
Monte Carlo estimation of the cumulative distribution functions of the *P*-values obtained by CpelTdm.jl when performing *randomization testing* using the differential test statistics *T*_MML_, *T*_NME_, and *T*_PDM_. A simulated *unmatched pairs group comparison* is considered within a small genomic region containing 4 CpG sites when the test and reference group contain 5 data samples each (*L* = 252 < 1000). (a) Region exhibits no differences (test: *α* = 0.5, *β* = 0.5; reference: *α* = 0.5, *β* = 0.5). (b) Region exhibits MML and PDM differences (test: *α* = 0.5, *β* = 0.5; reference: *α* = −0.5, *β* = 0.5). (c) Region exhibits NME and PDM differences (test: *α* = 0.0, *β* = 0.5; reference: *α* = 0, *β* = 0). (d) Region exhibits MML, NME, and PDM differences (test: *α* = 0.5, *β* = 0.5; reference: *α* = 0, *β* = 0). The marked (red stars) cumulative distribution functions indicate that CpelTdm.jl detects statistically significant differences with a rate that is no less than 95% (i.e., Pr[*p* ≤ 0.05] ≥ 0.95), in agreement with the region’s known differential behavior. In the remaining cases, Pr[*p* ≤ 0.05] = 0.05, indicating a 5% probability of Type I error.

**Figure 5.**
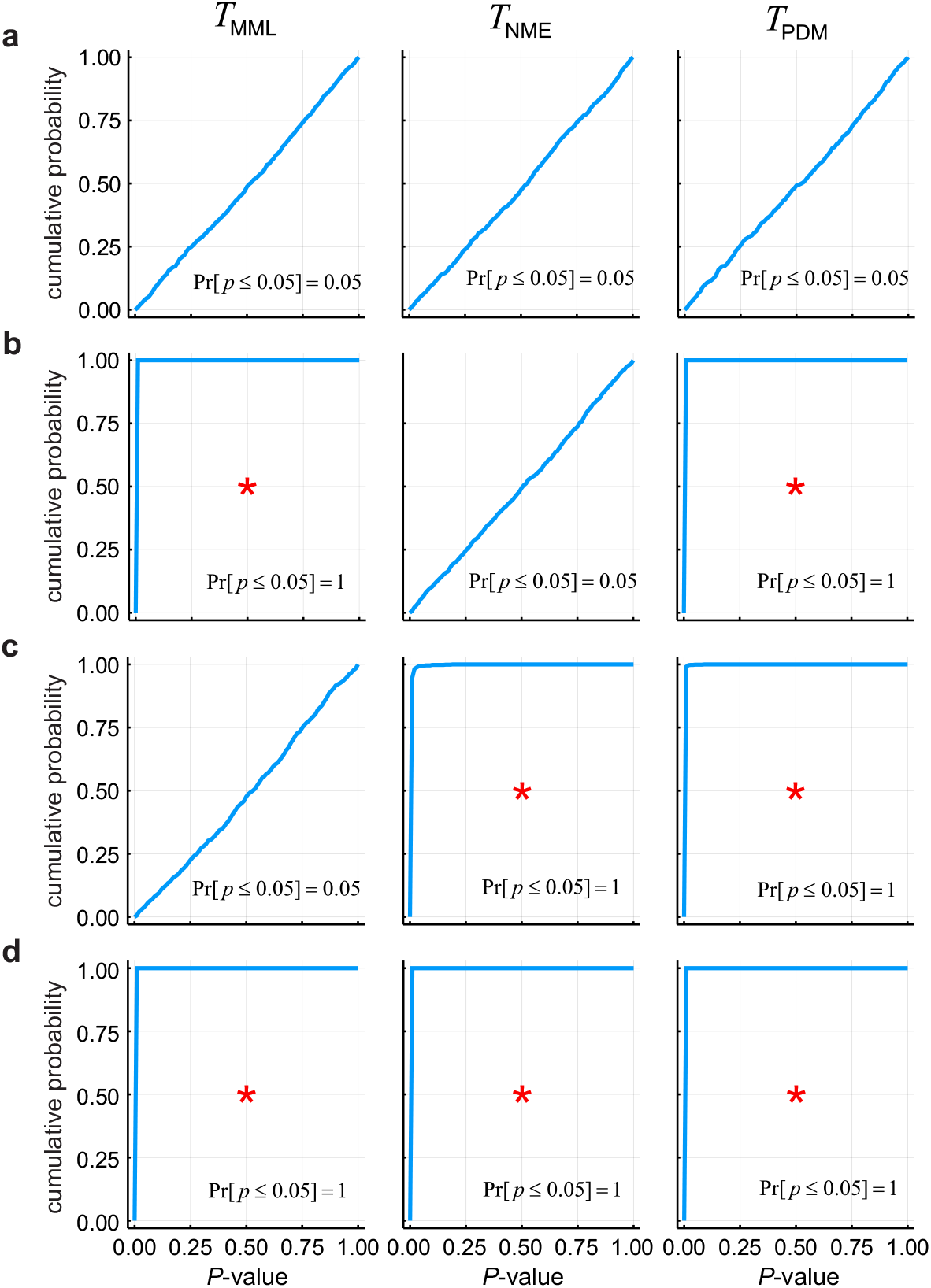
Monte Carlo estimation of the cumulative distribution functions of the *P*-values obtained by CpelTdm.jl when performing *Monte Carlo based permutation testing* using the differential test statistics *T*_MML_, *T*_NME_, and *T*_PDM_. A simulated *unmatched pairs group comparison* is considered within a small genomic region containing 4 CpG sites when the test and reference group contain 7 data samples each (*L* = 3432 > 1000). (a) Region exhibits no differences (test: *α* = 0.5, *β* = 0.5; reference: *α* = 0.5, *β* = 0.5). (b) Region exhibits MML and PDM differences (test: *α* = 0.5, *β* = 0.5; reference: *α* = −0.5, *β* = 0.5). (c) Region exhibits NME and PDM differences (test: *α* = 0.0, *β* = 0.5; reference: *α* = 0, *β* = 0). (d) Region exhibits MML, NME, and PDM differences (test: *α* = 0.5, *β* = 0.5; reference: *α* = 0, *β* = 0). The marked (red stars) cumulative distribution functions indicate that CpelTdm.jl detects statistically significant differences 100% of the time (i.e., Pr[*p* ≤ 0.05] = 1), in agreement with the region’s known differential behavior. In the remaining cases, Pr[*p* ≤ 0.05] = 0.05, indicating a 5% probability of Type I error.

**Figure 6.**
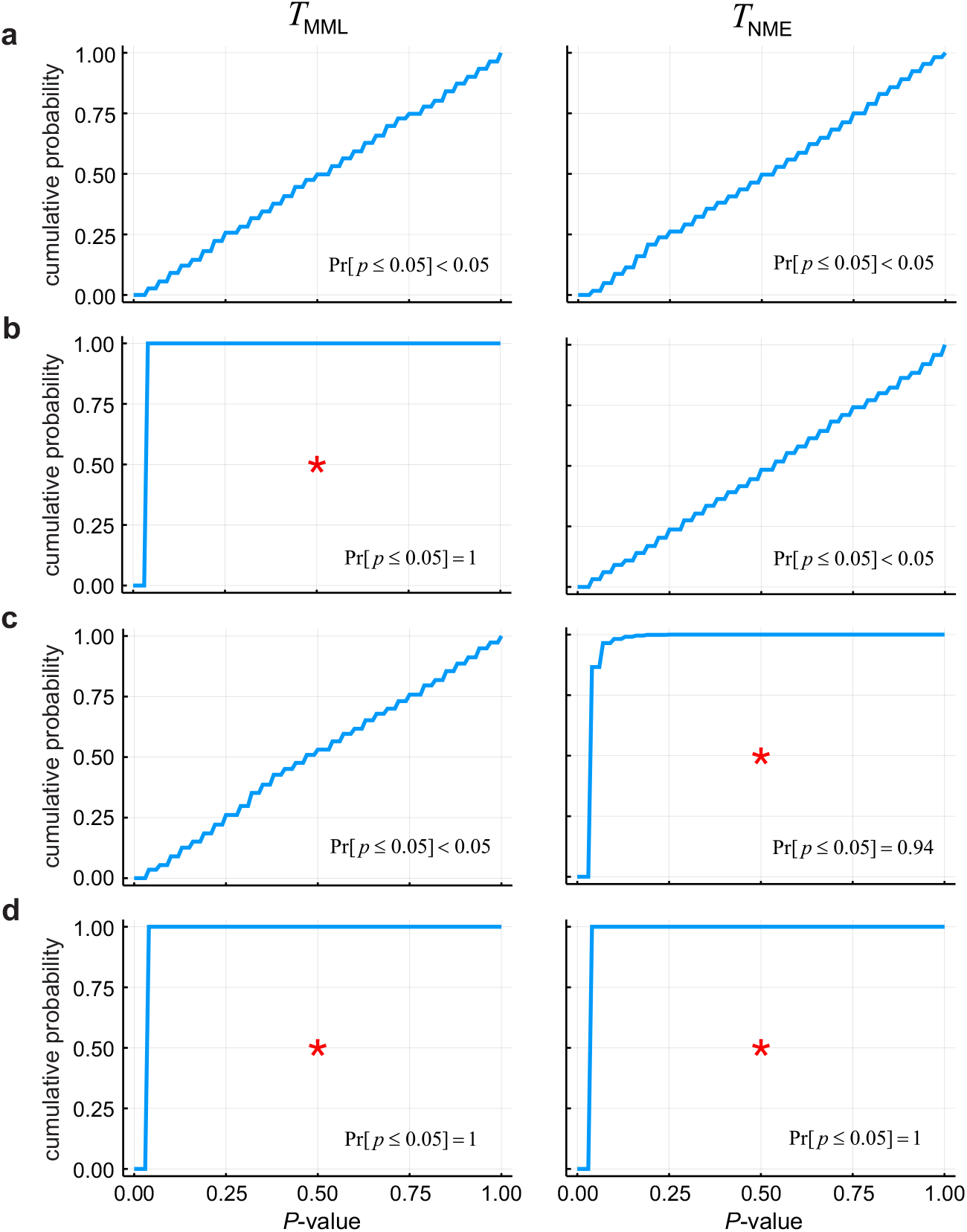
Monte Carlo estimation of the cumulative distribution functions of the *P*-values obtained by CpelTdm.jl when performing *randomization testing* using the differential test statistics *T*_MML_ and *T*_NME_. A simulated *matched pairs group comparison* is considered within a small genomic region containing 4 CpG sites when the test and reference group contain 6 data samples each (*L* = 64 < 1000). (a) Region exhibits no differences (test: *α* = 0.5, *β* = 0.5; reference: *α* = 0.5, *β* = 0.5). (b) Region exhibits MML and PDM differences (test: *α* = 0.5, *β* = 0.5; reference: *α* = −0.5, *β* = 0.5). (c) Region exhibits NME and PDM differences (test: *α* = 0.0, *β* = 0.5; reference: *α* = 0, *β* = 0). (d) Region exhibits MML, NME, and PDM differences (test: *α* = 0.5, *β* = 0.5; reference: *α* = 0, *β* = 0). The marked (red stars) cumulative distribution functions indicate that CpelTdm.jl detects statistically significant differences with a rate that is no less than 95% (i.e., Pr[*p* ≤ 0.05] ≥ 0.95), in agreement with the region’s known differential behavior. In the remaining cases, Pr[*p* ≤ 0.05] = 0.05, indicating a 5% probability of Type I error.

**Figure 7.**
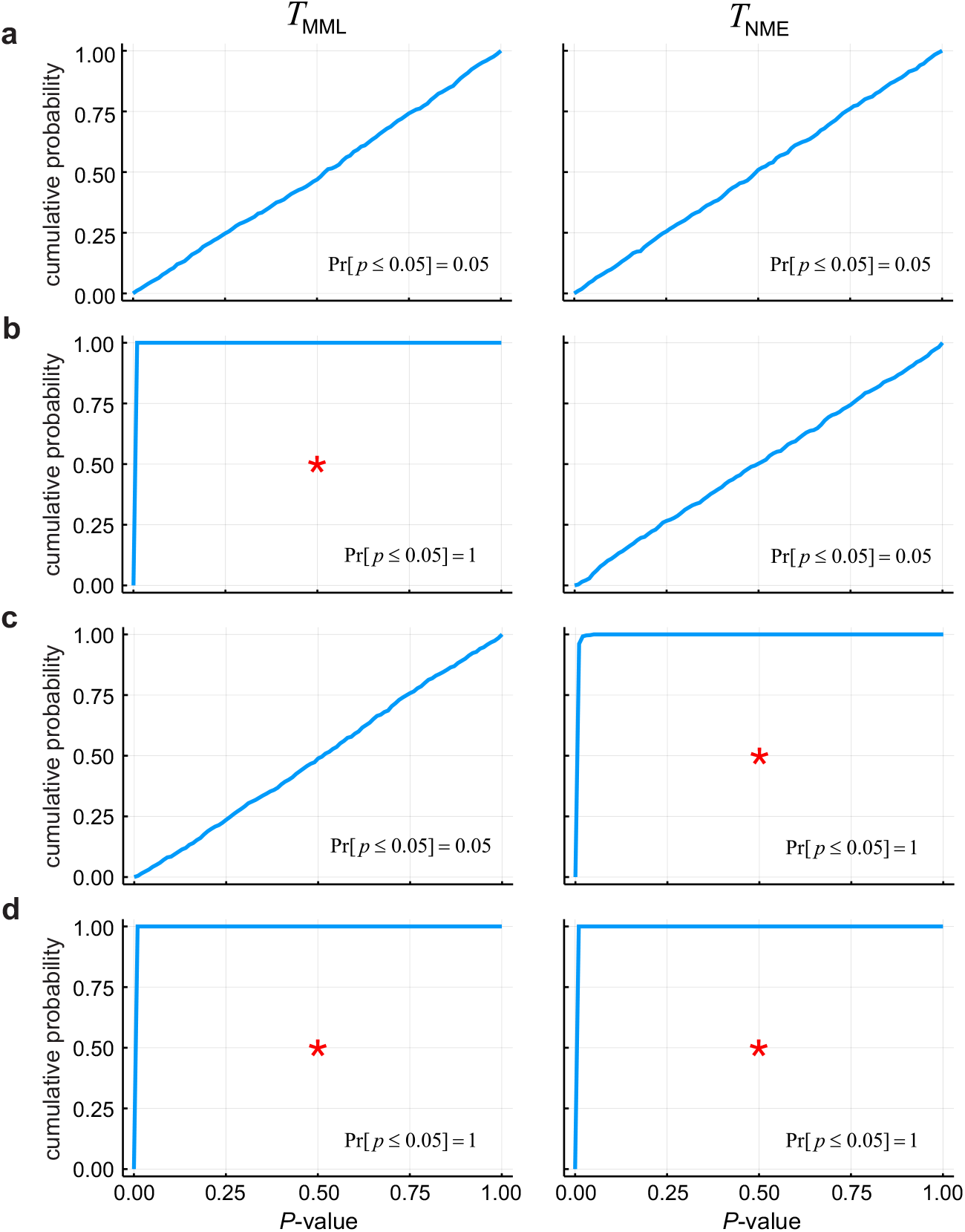
Monte Carlo estimation of the cumulative distribution functions of the *P*-values obtained by CpelTdm.jl when performing *Monte Carlo based permutation testing* using the differential test statistics *T*_MML_ and *T*_NME_. A simulated *matched pairs group comparison* is considered within a small genomic region containing 4 CpG sites when the test and reference group contain 10 data samples each (*L* = 1024 > 1000). (a) Region exhibits no differences (test: *α* = 0.5, *β* = 0.5; reference: *α* = 0.5, *β* = 0.5). (b) Region exhibits MML and PDM differences (test: *α* = 0.5, *β* = 0.5; reference: *α* = −0.5, *β* = 0.5). (c) Region exhibits NME and PDM differences (test: *α* = 0.0, *β* = 0.5; reference: *α* = 0, *β* = 0). (d) Region exhibits MML, NME, and PDM differences (test: *α* = 0.5, *β* = 0.5; reference: *α* = 0, *β* = 0). The marked (red stars) cumulative distribution functions indicate that CpelTdm.jl detects statistically significant differences 100% of the time (i.e., Pr[*p* ≤ 0.05] = 1), in agreement with the region’s known differential behavior. In the remaining cases, Pr[*p* ≤ 0.05] = 0.05, indicating a 5% probability of Type I error.

We also performed an unmatched-pair comparison over CpG islands (CGIs) between a group of 7 acute promyelocytic leukemia (APL) samples and a group of 6 remission samples after treatment using a previously published RRBS dataset; see Table 1. Densities of MML and NME values demonstrated an overall reduction of methylation level in the remission samples, consistently with previous results (Hebestreit *et al*., 2013), as well as a reduction in methylation entropy leading to a more ordered methylation landscape after treatment; see Fig. 8. This behavior was also reflected by the distributions of differential MML and NME test statistic values and was associated with differences in PDM; see Fig. 9a. Q-Q plots obtained by differential analysis for each test statistic revealed that most observed *P*-values below 0.05 were much smaller that the corresponding expected *P*-values under the null hypothesis, providing strong evidence of the existence of methylation differences between the two groups; see Fig. 9b. Multiple hypothesis testing identified a number of CGIs exhibiting significant differences in MML, NME, and PDM (*Q*-value ≤ 0.1), which were linked to the promoter regions of 618 genes; see Fig. 10 and Table 2. Interestingly, gene set enrichment analysis (GSEA) of the list of genes exhibiting significant PDM differences at their promoter regions revealed enrichments related to development, differentiation, morphogenesis, signaling, transcription, and chromatin/chromosome formation and organization (see Table 3), pointing to their importance in APL.

**Figure 8.**
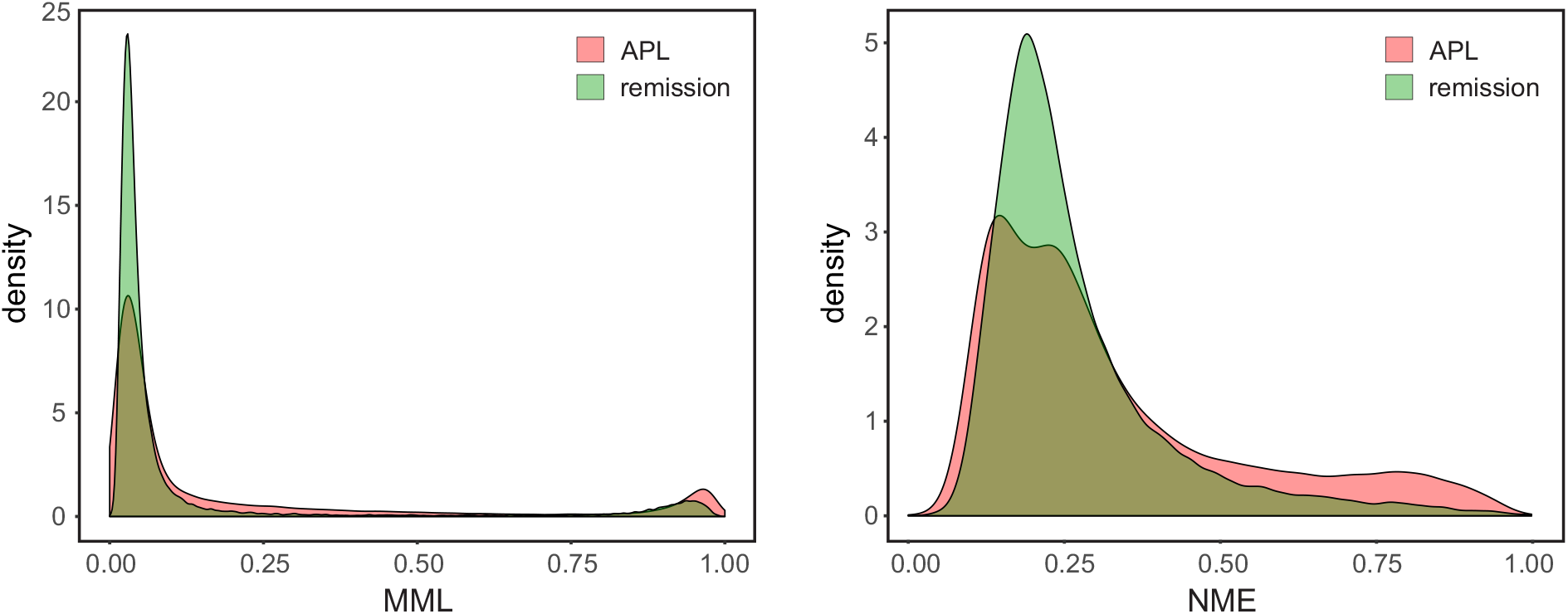
Densities of mean methylation level (MML) and normalized methylation entropy (NME) values within CGI analysis regions computed by CpelTdm.jl in an unmatched pairs group comparison between acute promyelocytic leukemia (APL) and remission samples. Notably, treatment results in an overall reduction of methylation level and methylation entropy leading to a more ordered methylation landscape after treatment.

**Figure 9.**
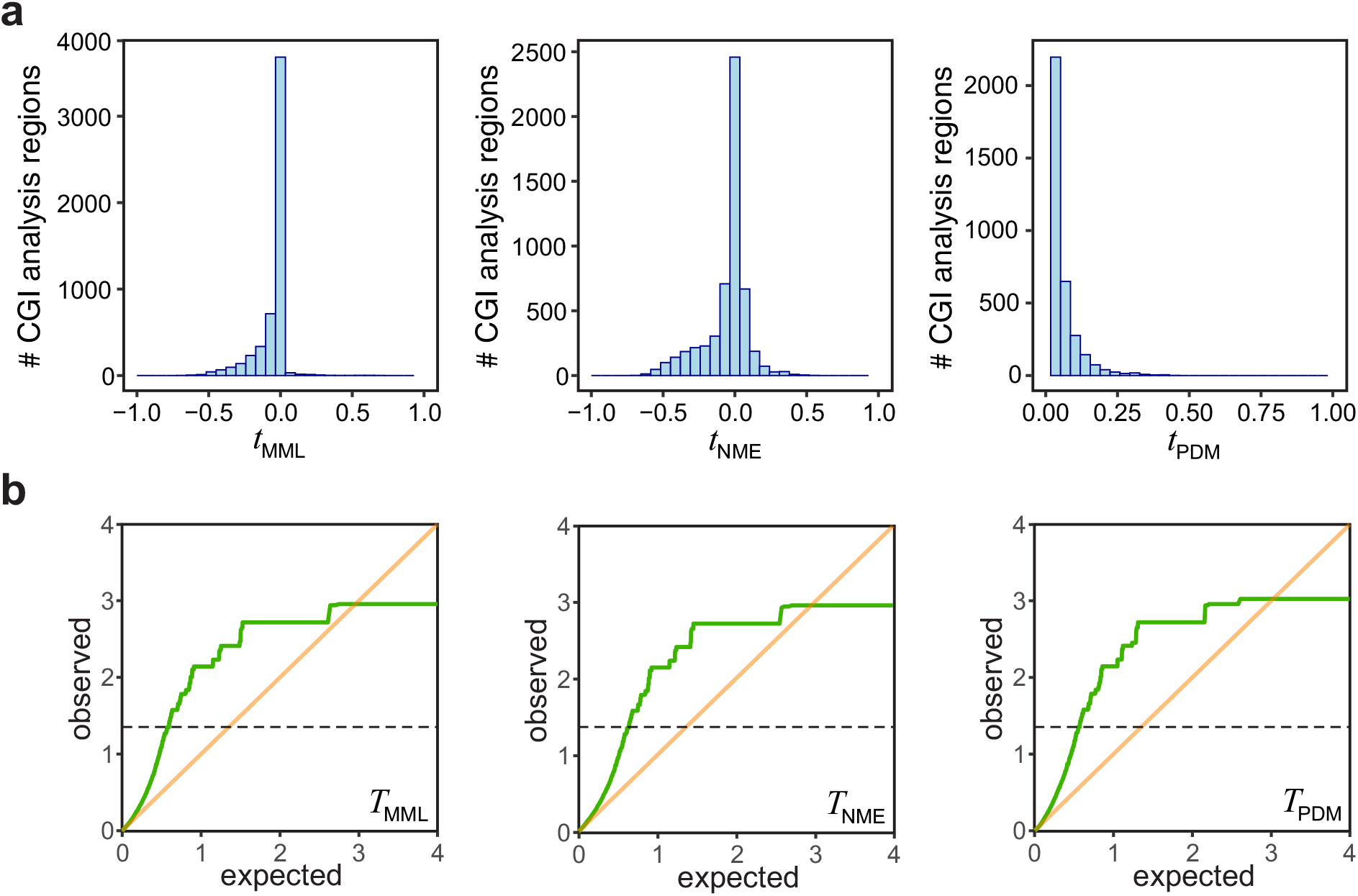
(a) Distributions of the differential MML, NME, and PDM test statistic values *t*_MML_, *t*_NME_, and *t*_PDM_ obtained by CpelTdm.jl from 5398 CGI analysis regions in the unmatched pairs group comparison between acute promyelocytic leukemia and remission samples. (b) Q-Q plots for each differential test statistic (green lines) of quantiles of – log_10_ *P*-values observed within the CGI analysis regions versus quantiles of expected – log_10_ *P*-values under the null hypothesis. Notably, since most observed *P*-values below 0.05 (over the dotted line) are located above the diagonal, they are much smaller than the corresponding *P*-values expected under the null hypothesis, thus providing strong evidence of the existence of meaningful differences in methylation stochasticity among the two groups.

**Figure 10.**
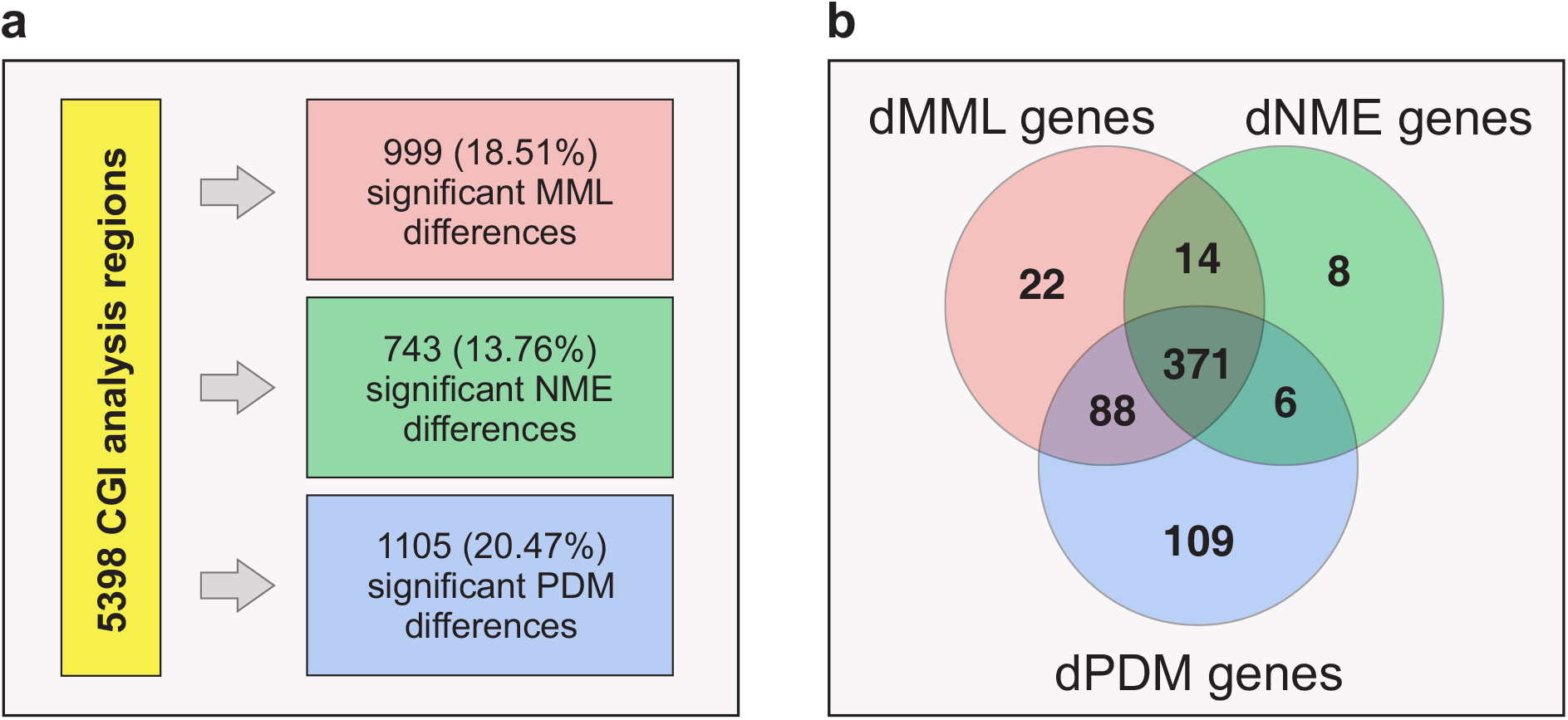
(a) Multiple hypothesis testing performed by CpelTdm.jl in the unmatched pairs group comparison between acute promyelocytic leukemia and remission samples identified CGI analysis regions with significant MML, NME, and PDM differences (*Q*-values ≤ 0.1). Notably, an unmatched pairs remission/remission group comparison did not find statistically significant analysis regions, as expected. (b) A total of 618 genes were identified to exhibit statistically significant MML, NME, or PDM differences within at least one CGI analysis region overlapping their promoter regions (4-kb window centered at TSS). Notably, 495 genes (80%) were associated with significant MML differences (dMML genes), 399 (65%) with significant NME differences (dNME genes), and 574 (93%) with significant PDM differences (dPDM genes). In addition, 22 genes were associated with only significant MML differences (e.g., the homeobox gene *HOXD8* implicated in stem cell differentiation and cancer, including APL, the proto-oncogene *RET* involved in activating signaling pathways that play a role in cell differentiation, growth, migration and survival, and the transcription factor *TCF7L1* implicated in gastric and colorectal cancers as well as in acute lymphoblastic leukemia). Moreover, 8 genes were associated with only significant NME differences (e.g., *IGFBP5* that is often dysregulated in cancer, *LAPTM4B*, an oncogene that has been found to be highly expressed in various tumors, and *PAWR*, a tumor suppressor that induces apoptosis in cancer cells). Finally, 109 genes were associated with only significant PDM differences (e.g., *AK1* and *SYNE2*, both implicated in acute myeloid leukemia, *ITPKA*, which is expressed in a broad range of tumor types but shows limited expression in normal cells, and *MPPED2* that plays a carcinogenetic role across different cancer types).

**Table 1.** Summary of the RRBS dataset used for targeted differential methylation analysis, which includes 7 acute promyelocytic leukemia (APL) samples and 6 remission samples (NCBI GEO accession number GSE42044).

| **Name** | **Sample** | **Condition** | **Sex** |
| --- | --- | --- | --- |
| apl1 | APL_10759 | APL | Female |
| apl2 | APL_7717 | APL | Female |
| apl3 | APL_10545 | APL | Male |
| apl4 | APL_19 | APL | Male |
| apl5 | APL_9882 | APL | Male |
| apl6 | APL_5 | APL | Female |
| apl7 | APL_10892 | APL | Female |
| remission1 | APL_10961 | Remission | Female |
| remission2 | APL_11624 | Remission | Female |
| remission3 | APL_11523 | Remission | Male |
| remission4 | APL_16 | Remission | Male |
| remission5 | APL_10325 | Remission | Male |
| remission6 | APL_5894 | Remission | Male |

**Table 2.**
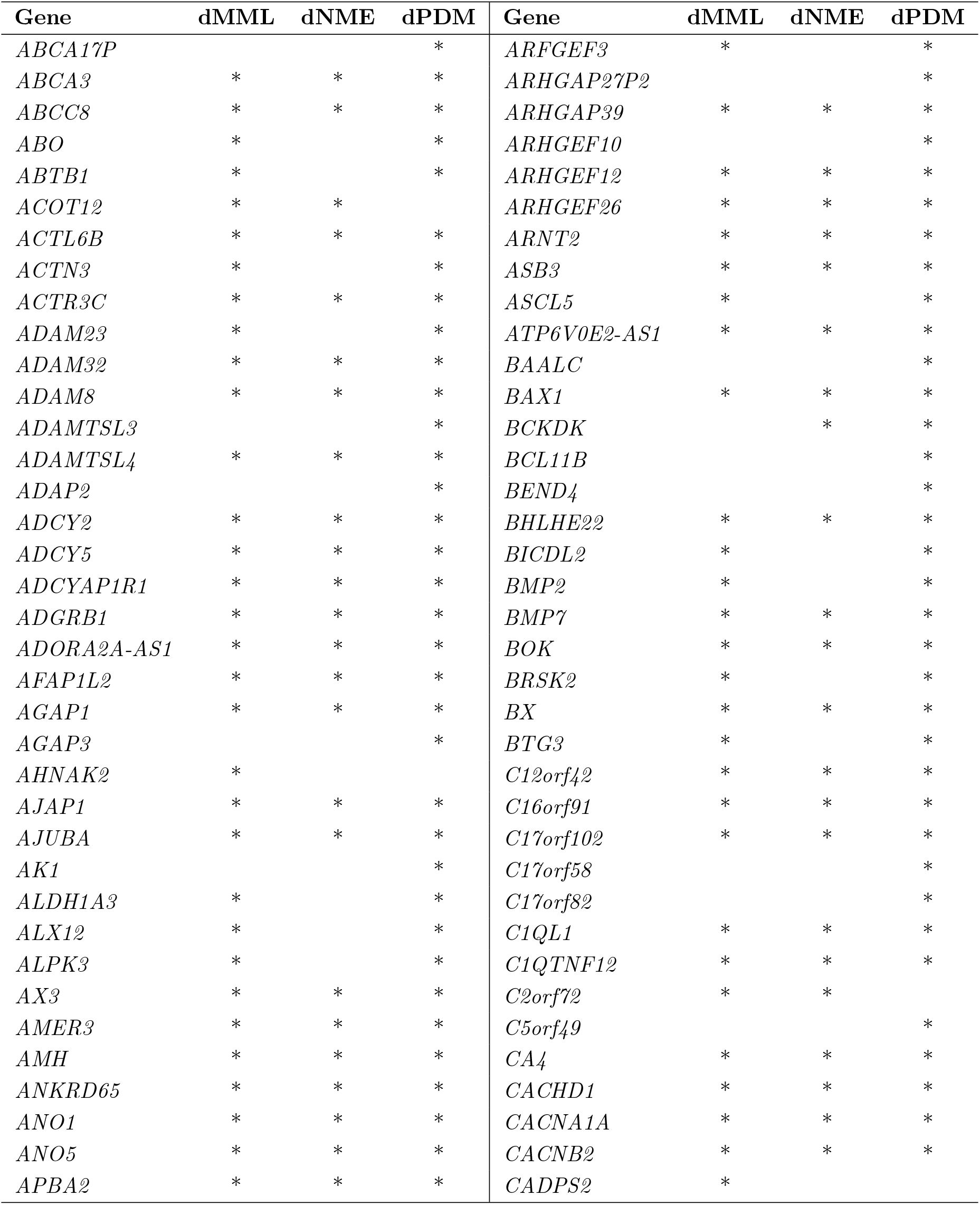
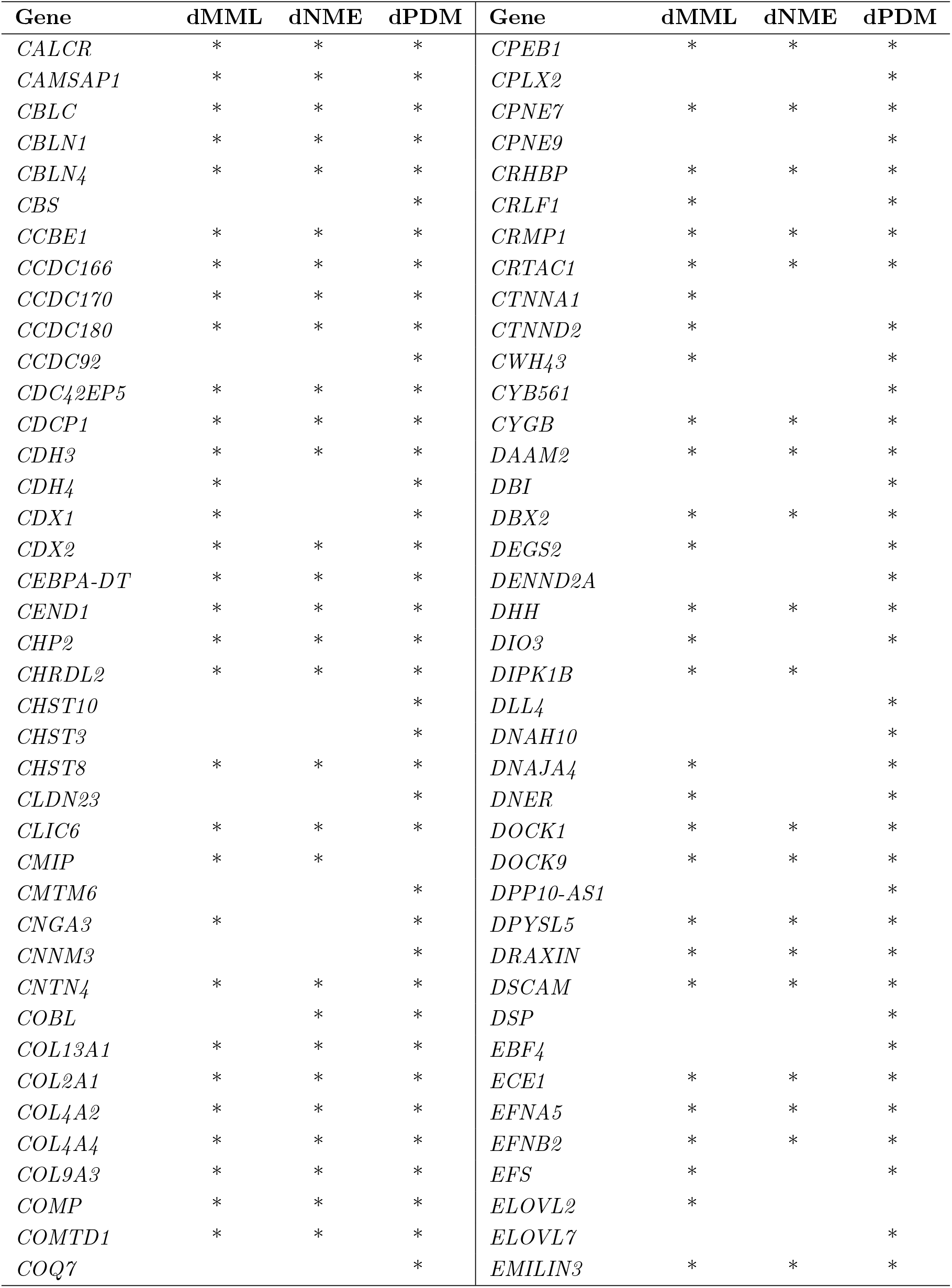
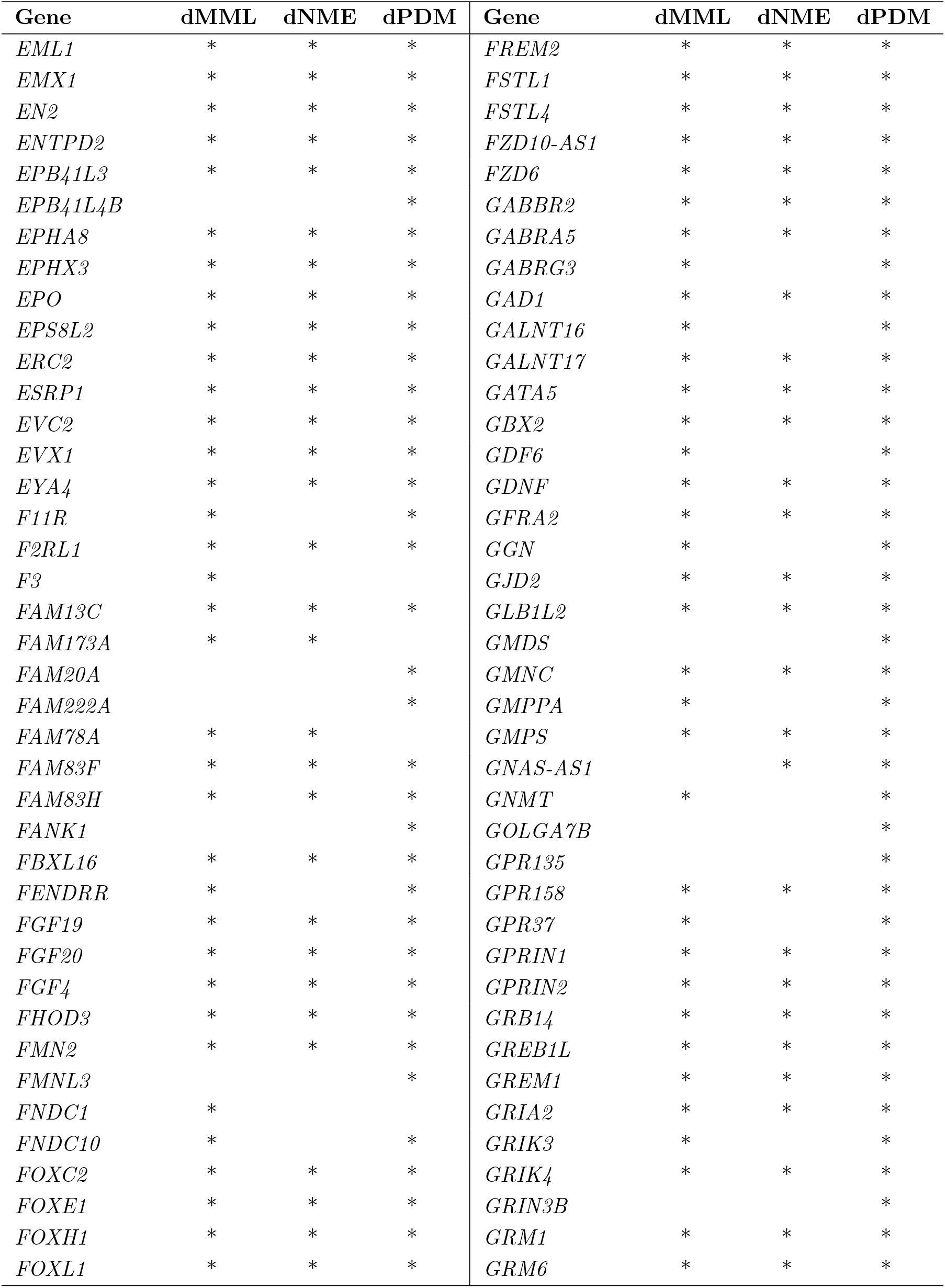
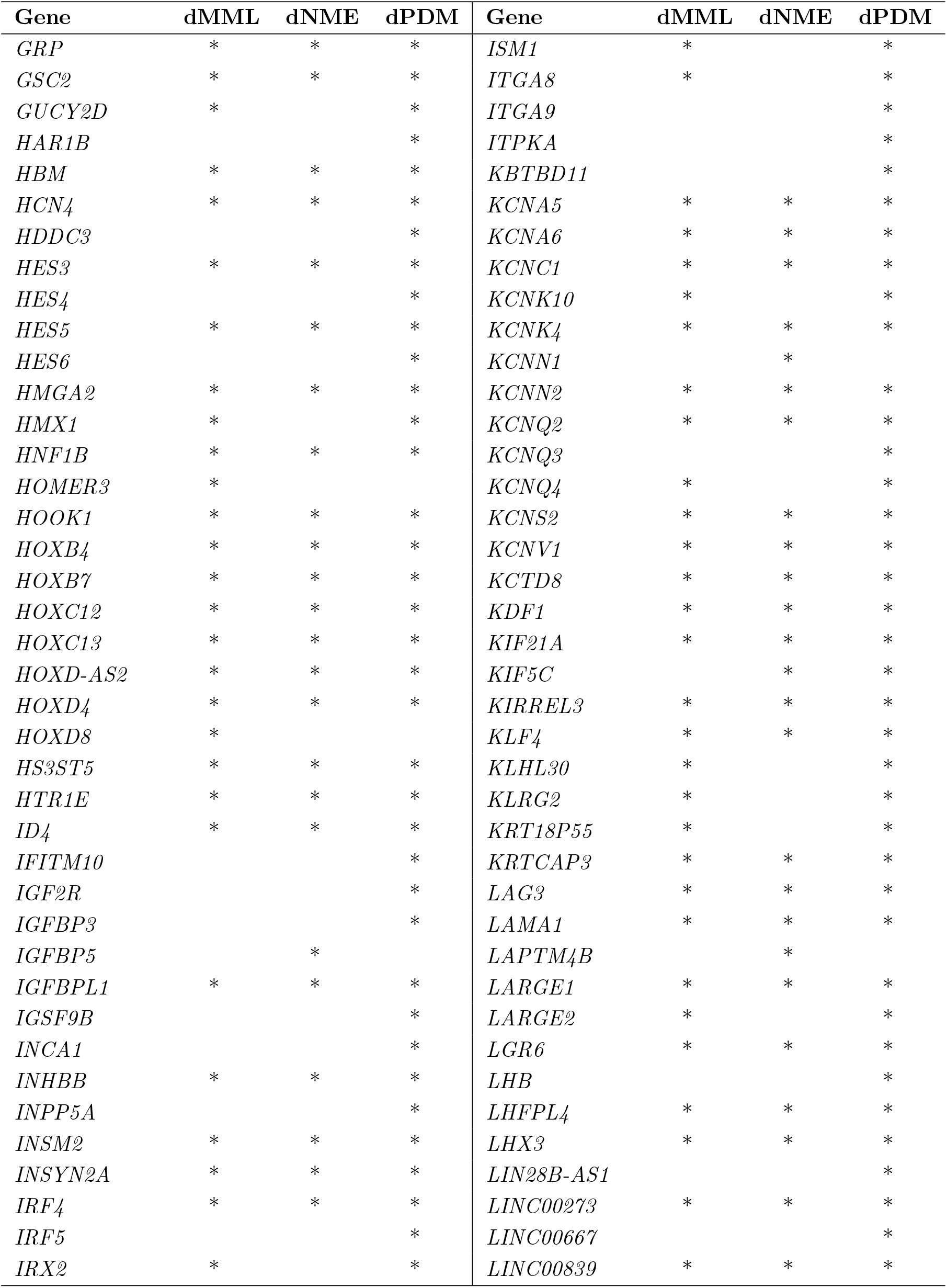
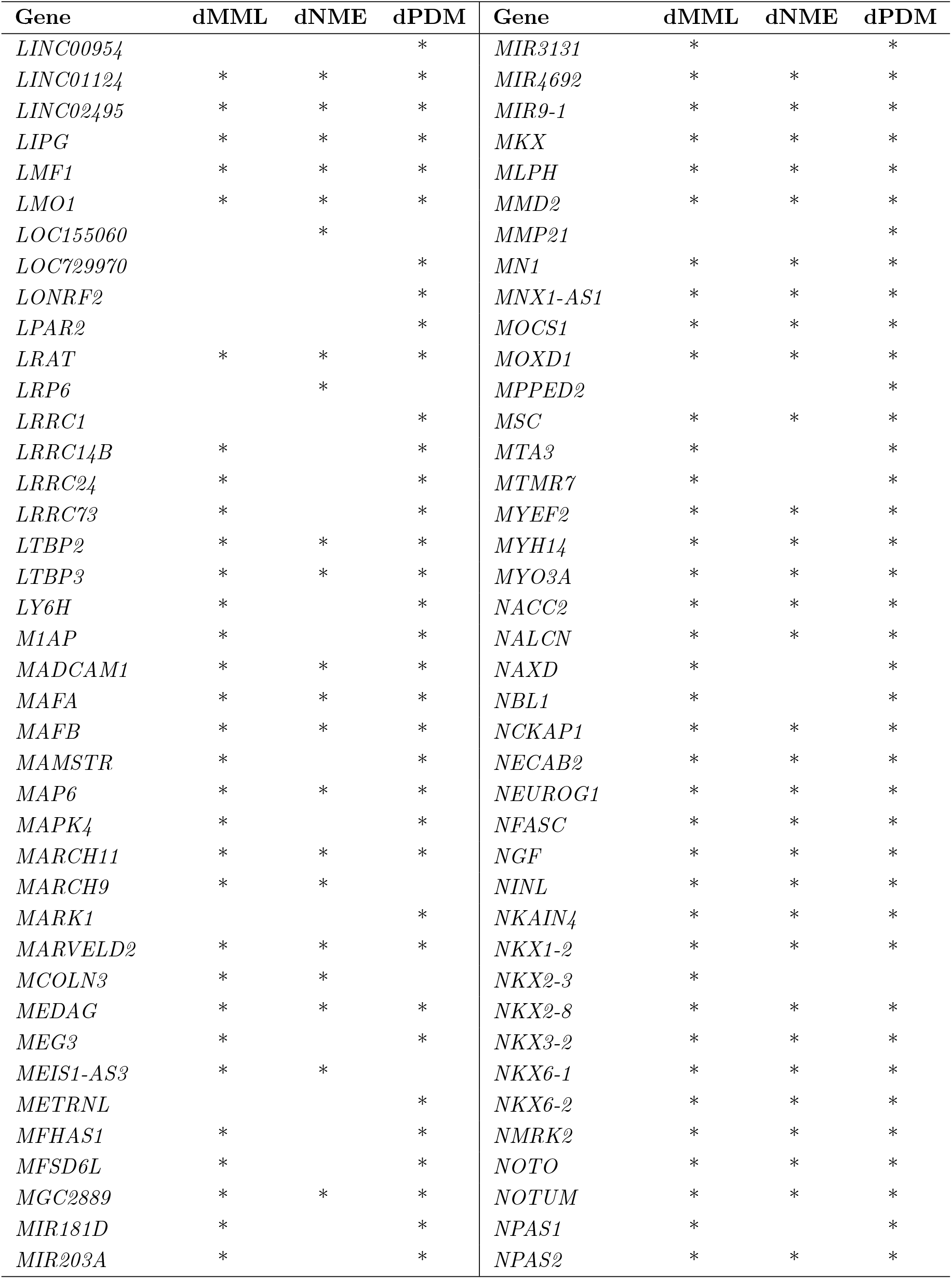
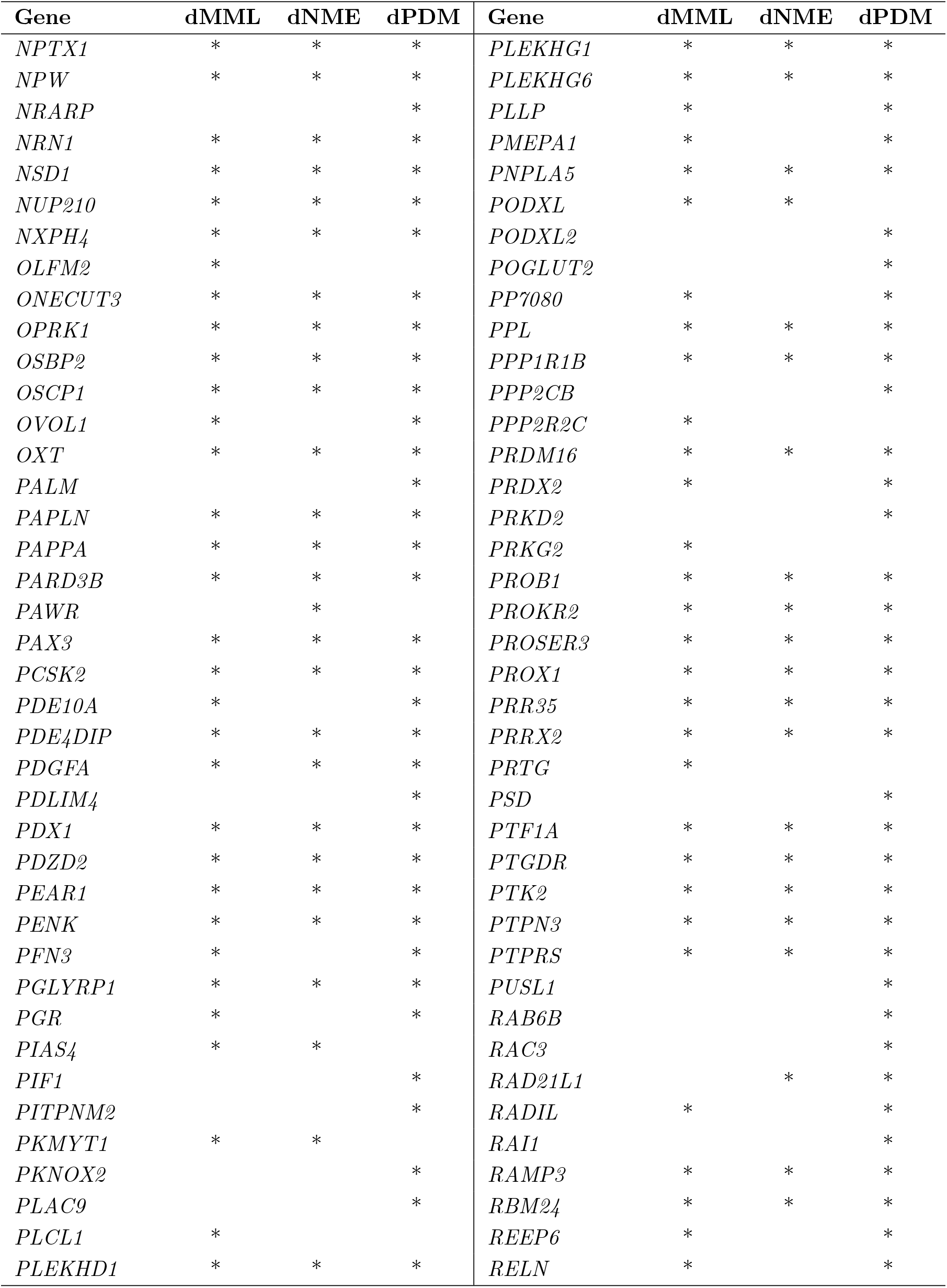
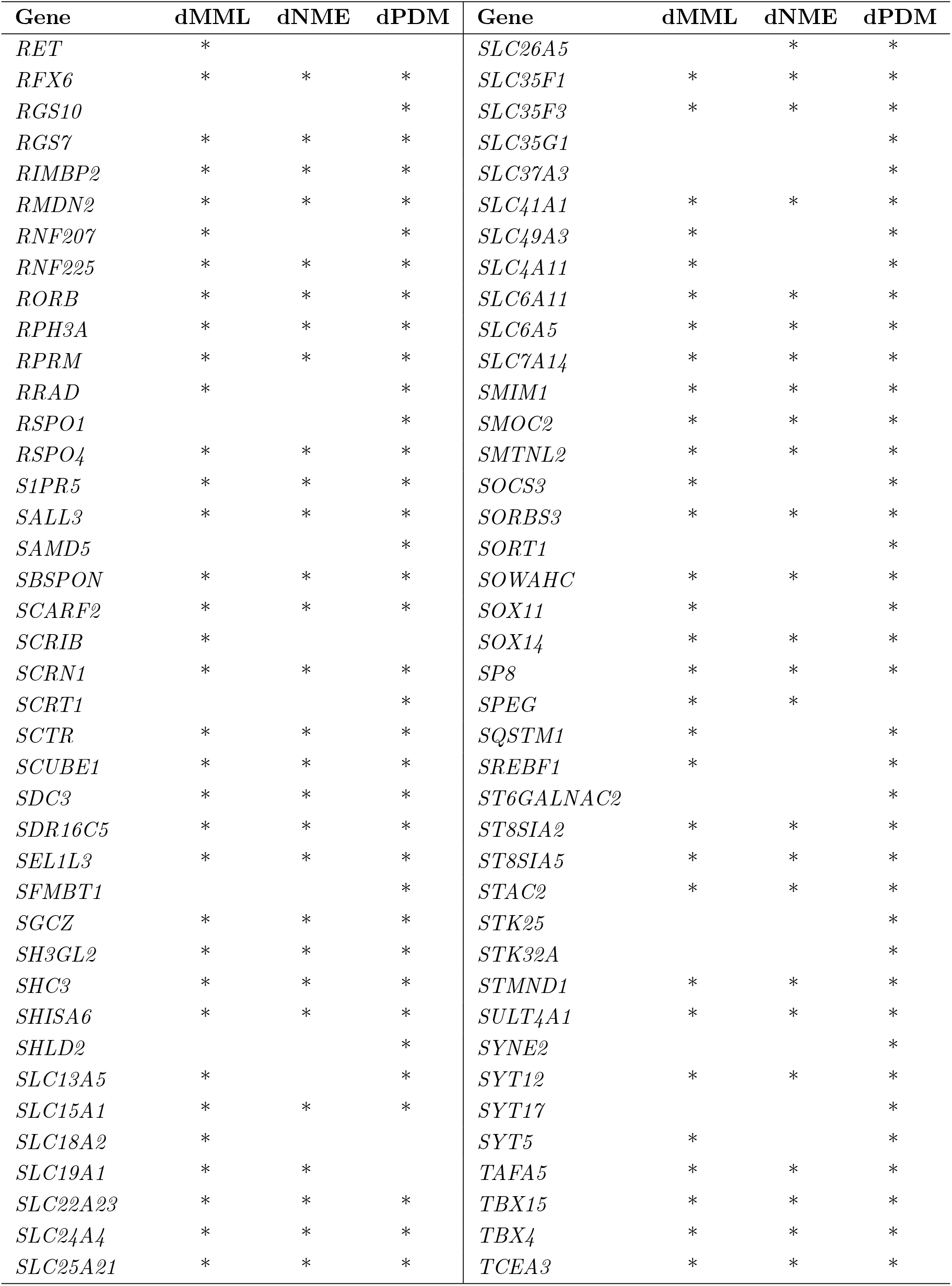
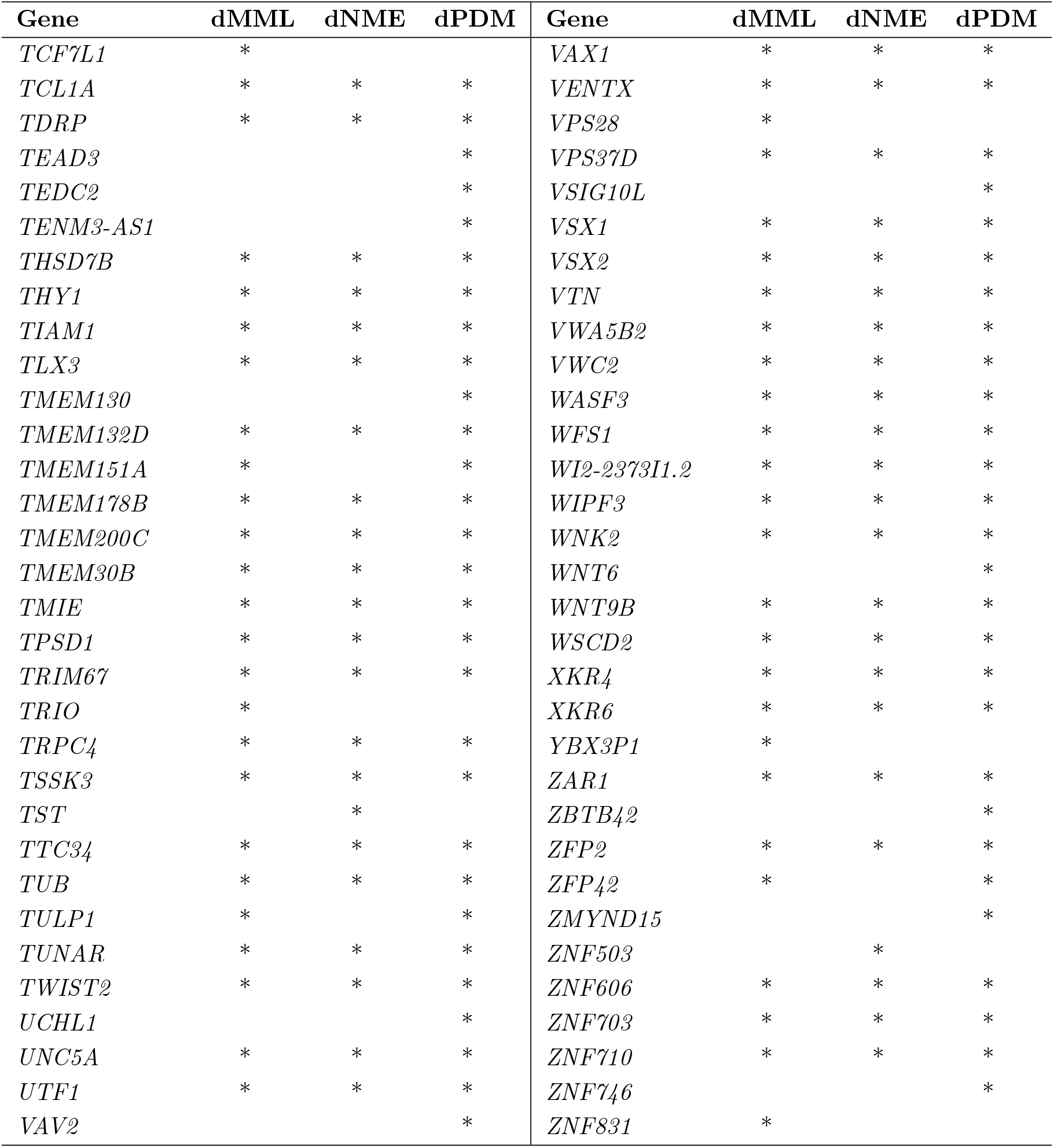
Genes identified by CpelTdm.jl to exhibit significant MML, NME, or PDM differences (dMML, dNME, and dPDM genes) within at least one CGI analysis region overlapping their promoter regions (4-kb window centered at TSS) in the unmatched pairs group comparison between the acute promyelocytic leukemia and remission samples.

**Table 3.** Gene set enrichment analysis (GSEA) results (overlaps with the ‘GO gene sets’ in MSigDB) of the 574 genes in Table 2 exhibiting statistically significant PDM differences (dPDM genes). GSEA Collection: C5; overlaps shown: 25; # genes in collection: 10,192; # genes in comparison: 570; # genes in universe: 38,404.

| Gene set name | Genes in<br>gene set | Genes in<br>overlap | FDR<br>Q-value |
| --- | --- | --- | --- |
| GO_NEUROGENESIS | 1625 | 108 | 7.62E-36 |
| GO_NEURON_DIFFERENTIATION | 1365 | 93 | 2.57E-31 |
| GO_CELL_CELL_SIGNALING | 1665 | 98 | 3.07E-28 |
| GO_DNA_BINDING_TRANSCRIPTION_FACTOR_ACTIVITY_RNA_POLYMERASE_II_SPECIFIC | 1094 | 79 | 5.28E-28 |
| GO_SEQUENCE_SPECIFIC_DNA_BINDING | 1189 | 79 | 1.11E-25 |
| GO_NEURON_PROJECTION | 1317 | 83 | 1.22E-25 |
| GO_DNA_BINDING_TRANSCRIPTION_FACTOR_ACTIVITY | 1334 | 81 | 6.14E-24 |
| GO_CHROMATIN | 1226 | 77 | 1.37E-23 |
| GO_TRANSCRIPTION_REGULATOR_ACTIVITY | 1718 | 90 | 1.85E-22 |
| GO_NUCLEAR_CHROMOSOME | 1273 | 76 | 5.61E-22 |
| GO_ANIMAL_ORGAN_MORPHOGENESIS | 1071 | 69 | 1.27E-21 |
| GO_REGULATION_OF_NERVOUS_SYSTEM_DEVELOPMENT | 928 | 62 | 4.89E-20 |
| GO_NEURON_DEVELOPMENT | 1114 | 68 | 4.89E-20 |
| GO_SYNAPSE | 1332 | 74 | 1.27E-19 |
| GO_REGULATORY_REGION_NUCLEIC_ACID_BINDING | 960 | 62 | 2.24E-19 |
| GO_REGULATION_OF_CELL_DIFFERENTIATION | 1881 | 88 | 7.21E-19 |
| GO_SOMATODENDRITIC_COMPARTMENT | 834 | 57 | 7.21E-19 |
| GO_CELLULAR_COMPONENT_MORPHOGENESIS | 1141 | 66 | 2.81E-18 |
| GO_CENTRAL_NERVOUS_SYSTEM_DEVELOPMENT | 995 | 61 | 5.29E-18 |
| GO_POSITIVE_REGULATION_OF_TRANSCRIPTION_BY_RNA_POLYMERASE_II | 1198 | 67 | 7.46E-18 |
| GO_PATTERN_SPECIFICATION_PROCESS | 446 | 41 | 9.31E-18 |
| GO_SEQUENCE_SPECIFIC_DOUBLE_STRANDED_DNA_BINDING | 889 | 57 | 1.15E-17 |
| GO_CELL_PROJECTION_ORGANIZATION | 1545 | 76 | 2.45E-17 |
| GO_CHROMOSOME | 1730 | 81 | 2.45E-17 |
| GO_EMBRYONIC_MORPHOGENESIS | 592 | 46 | 3.16E-17 |

The previous APL analysis took about 7 hours using 22 CPUs and about 1 GB of RAM per CPU. The required input CGI BED file and the output bedGraph files can be freely downloaded from http://www.cis.jhu.edu/~goutsias/data/CpelTdm_output.zip.

## Methods

### Defining analysis regions

CpelTdm.jl (see Fig. 1 for a flowchart) performs targeted differential DNA methylation analysis in order to identify statistically significant differences in methylation stochasticity between two groups of samples within genomic regions of interest that may include CpG islands (CGIs), promoter regions, enhancers, CpG clusters computed by the method proposed by Hebestreit *et al*. (2013), etc. These regions must be specified by the user in a single-track input BED file (see Fig. 1). To strike a balance between computational cost and estimation performance, CpelTdm.jl partitions any connected target genomic region whose size is more than 1-kb into a minimum number of equally-sized subregions of size ≤ 1-kb. It then independently performs downstream methylation analysis only within subregions for which the coverage at each CpG site is at least 5, which we refer to as ‘analysis regions’.

### Modeling DNA methylation stochasticity

CpelTdm.jl models methylation stochasticity within an analysis region 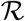 that contains *N* CpG sites *n* = 1,2,…, *N* using the probability distribution of methylation (PDM) *π*(x) = Pr[**X** = **x**] of an *N* × 1 random vector **X** = [*X*_1_ *X*_2_ ⋯ *X_N_*]^*T*^, where *X_n_* = 0 if the *n*-th CpG site is unmethylated and *X_n_* = 1 if methylated. It then partitions the analysis region 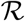 into the minimum number of equally-sized non-overlapping subregions 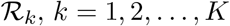, such that the size of each subregion does not exceed a maximum size *G* (in bp) and characterizes the random methylation state **X** by means of a previously introduced correlated potential energy landscape (CPEL) model, given by (Abante *et al*., 2020)

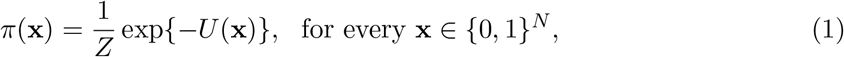

with potential energy function

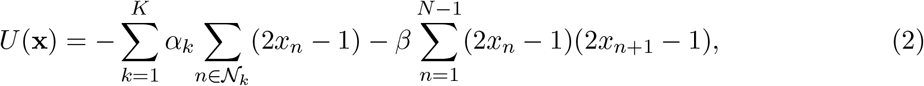

and partition function

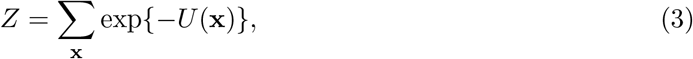

where 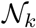 denotes the set of all CpG sites within 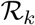, *α_k_* is a parameter that influences the propensity of CpG sites in the *k*-th subregion 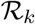 to be methylated due to non-cooperative factors, and *β* is a parameter that accounts for the fact that the methylation status of two contiguous CpG sites *n* and *n* + 1 within the entire analysis region 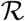 could be correlated. Notably, parameter *G* determines the ‘granularity’ of modeling with its value chosen in a way that strikes a balance between the scale of differential methylation analysis and the success rate of estimating the parameters of the CPEL model from available bisulfite data. CpelTdm.jl uses a default granularity of *G* = 250 bp and performs necessary partition function computations and PDM marginalizations using a set of previously developed computationally efficient equations. These equations involve multiplications of 2 × 2 matrices determined by spectral decompositions requiring computation of eigenvalues and eigenvectors, which are performed analytically using exact formulas (Abante *et al*., 2020).

### Performing parameter estimation

Given *I* independent and possibly partially observed reads **x**_1_, **x**_1_,…,**x**_*I*_ of the methylation state in a given analysis region, CpelTdm.jl estimates the parameters ***θ*** = [*α*_1_ *α*_2_ ⋯ *α_K_ β*]^*T*^ of the potential energy function of the CPEL model (see Fig. 1) by solving the following maximum-likelihood optimization problem:

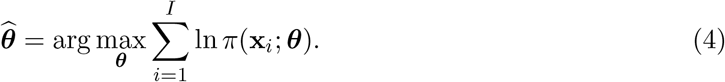

This is done by using Simulated Annealing with a temperature reduction factor of 10^−4^, a global optimization algorithm that was previously determined to yield superior estimation performance when compared to a number of alternative methods (Abante *et al*., 2020).

### Computing mean methylation levels

CpelTdm.jl quantifies the average amount of methylation within an analysis region containing *N* CpG sites using the mean methylation level (MML), given by (Jenkinson *et al*., 2017)

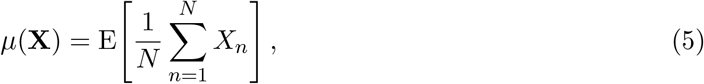

where E[·] denotes expectation. The MML evaluates the fraction of CpG sites that are methylated within an analysis region, taking its minimum value when all CpG sites are unmethylated and achieving its maximum value when all CpG sites are methylated; see Fig. 2. It can be shown that (Abante *et al*., 2020)

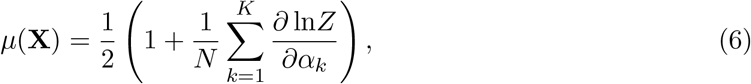

a formula employed by CpelTdm.jl to efficiently evaluate the MML by computing the derivatives of the logarithm of the partition function with respect to the parameters *α_k_* using a standard derivative approximation technique.

### Computing normalized methylation entropies

CpelTdm.jl quantifies the amount of methylation stochasticity (pattern heterogeneity) within an analysis region containing *N* CpG sites using the normalized methylation entropy (NME), given by (Jenkinson *et al*., 2017)

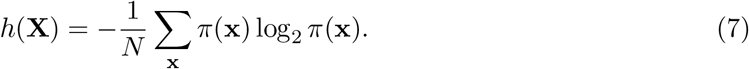

The NME ranges between 0 and 1, taking its minimum value when a single methylation pattern is observed within the analysis region (perfectly ordered methylation) and achieving its maximum value when all methylation patterns are equally likely (fully disordered methylation); see Fig. 2. It can be shown that (Abante *et al*., 2020)

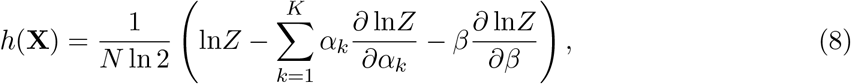

a formula used by CpelTdm.jl to efficiently evaluate the NME by employing the previously computed derivatives of the logarithm of the partition function with respect to the parameters *α_k_* and by also computing the derivative of the logarithm of the partition function with respect to parameter *β* using a standard derivative approximation technique.

### Computing coefficients of methylation divergence

Within an analysis region, CpelTdm.jl quantifies differences between the PDMs *π*_1_(**x**) and *π*_2_(**x**) of the methylation states **X**_1_ and **X**_2_ in a data sample from the first group and a data sample from the second group by using the coefficient of methylation divergence (CMD), which we define by

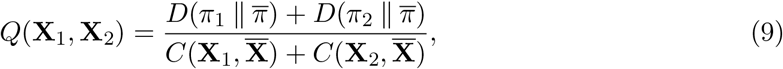

where 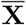 is a random vector characterized by a CPEL model 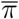 whose potential energy function is the average of the potential energy functions associated with **X**_1_ and **X**_2_,

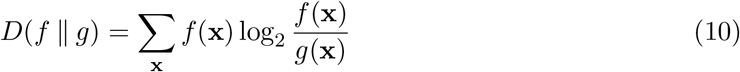

is the Kullback-Leibler (KL) divergence between two probability distributions *f* and *g*, and

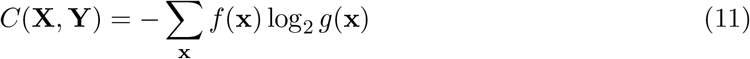

is the cross entropy between two random vectors **X** and **Y** characterized by probability distributions *f* and *g*, respectively. Notably, 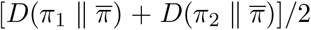 is known as the *geometric* Jensen-Shannon divergence (Nielsen, 2019), since it quantifies differences between the probability distributions *π*_1_ and *π*_2_ and their geometric average 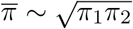, as compared to the standard Jensen-Shannon distance that quantifies differences between *π*_1_ and *π*_2_ and their arithmetic average (*π*_1_ + *π*_2_)/2. The CMD ranges between 0 and 1, taking its minimum value when the two PDMs *π*_1_ and *π*_2_ are identical and achieving its maximum when the supports of the probability distributions *π*_1_ and *π*_2_ do not overlap with the support of 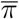, indicating radically different PDMs between the two groups.

It can be shown that

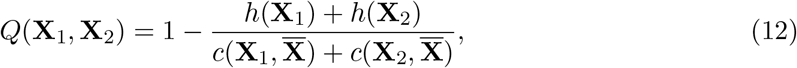

where *h* is the NME and *c* is the normalized cross methylation entropy (i.e., the cross entropy divided by *N*). Moreover,

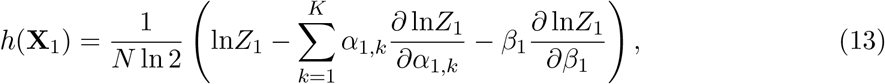

and

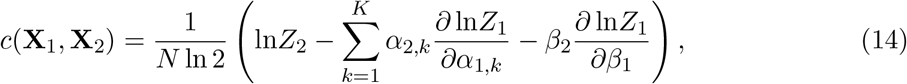

where {*α*_1,*k*_, *k* = 1,2,…,*K*, *β*_1_, *Z*_1_} and {*α*_2,*k*_, *k* = 1,2,…,*K*, *β*_2_, *Z*_2_} are respectively the energy parameters and partition functions of the CPEL models associated with **X**_1_ and **X**_2_. Using these formulas, CpelTdm.jl efficiently evaluates the CMD by computing partition function derivatives using a standard approximation technique.

### Performing hypothesis testing

CpelTdm.jl identifies regions of the genome that demonstrate statistically significant differences in DNA methylation stochasticity between two groups of samples by performing an unmatched or matched pairs group comparison. This is achieved by using permutation methods (Ernst, 2004) to test, for each analysis region, the null hypothesis *H*_0_ that each pair of samples exhibits no difference in methylation stochasticity (quantified by the MML, NME, or PDM) regardless of the specific group sample assignment.

#### Unmatched pairs group comparison

In an unmatched pairs comparison associated with a group of *M*_1_ values of a given statistical summary (MML, NME, or PDM) of methylation stochasticity within an analysis region and another group of *M*_2_ values associated with the same analysis region, CpelTdm.jl performs hypothesis testing using the following differential test statistics (see Fig. 1):

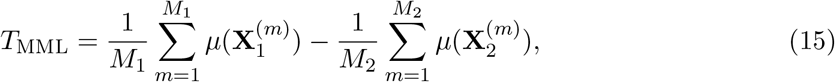

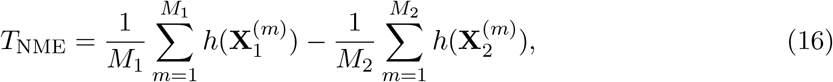

and

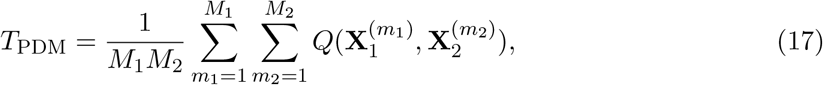

where 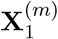 and 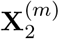 are the m-th methylation states of the analysis region within the first and second group, respectively. The test statistic *T*_MML_ quantifies the difference between the average of th mean methylation levels in each groups, *T*_NME_ assesses the difference between the averages of the normalized methylation entropies, and *T*_PDM_ quantifies the average of the PDM differences observed between the two groups; see Fig. 3.

For each test statistic *T*, a (two-tailed) hypothesis test requires knowledge of the null cumulative distribution function *F*_0_(*t*) = Pr[|*T*| < *t* |*H*_0_], which can then be used to calculate the *P*-value associated with an observation *t*\* of *T* by *p* = 1 – *F*_0_(|*t*\*|). To compute *F*_0_(*t*) for an analysis region, CpelTdm.jl uses a ‘randomization model’ that randomly assigns the available samples to the two groups, thus forming *L* = (*M*_1_ + *M*_2_)!/*M*_1_!*M*_2_! group assignments, provided that *L* is less than a maximum allowable number *L*_0_, which is taken to be *L*_0_ = 1000 by default; see Fig. 1. Under the null hypothesis, the group assignments are equally likely (with probability 1/*L*) to produce the expected outcome. As a consequence, the null cumulative distribution function *F*_0_(*t*) of a test statistic *T* will be given by 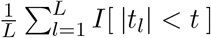, where *t_l_, l* = 1,2,…, *L*, are values of *T* computed for each group assignment and *I*[·] is the Iverson bracket, taking value 1 when its argument is true and 0 otherwise. This leads to an *exact P*-value computation given by (see Fig. 1)

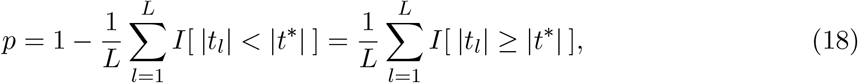

where *t*\* is the value of the *T* statistic calculated from the given data.

Note that the only possible values for *p* produced by the previous randomization testing approach are 1/*L*, 2/*L*,…, 1. Moreover, if the significance level of the test is set to *a* = *k*/*L* for some integer *k* such that *p* can take value *k*/*L*, then

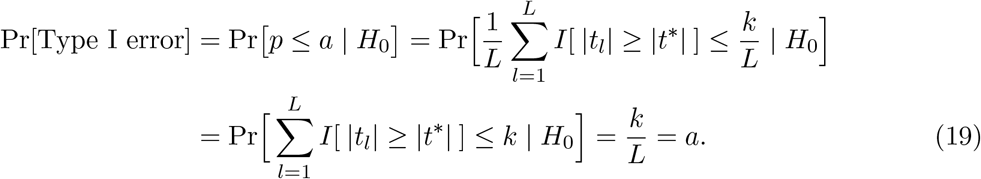

On the other hand, if *a* = *k*/*L* for some integer *k* such that *p* cannot take a value *k*/*L*, then Pr[Type I error] < *a* and the test will be conservative. As a consequence, the randomization testing procedure used by CpelTdm.jl can always control the Type I error (false positives).

When *L* ≥ *L*_0_, the previous method becomes computationally intensive. For this reason, CpelTdm.jl switches to a hypothesis testing approach that estimates the *P*-value using a permutation test based on Monte Carlo sampling; see Fig. 1. In this case, *L* distinct sample permutations are performed by assigning *M*_1_ samples to the first group and the remaining samples to the second group. The null cumulative distribution function is then approximated by independently sampling, *L*_0_ − 1 times, the set of *L* permutations with equal probability and by setting 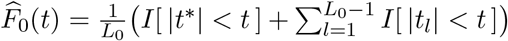, since this method produces *L*_0_ test statistic values, including the value *t*\* computed from the data. In this case, the *P*-value is approximated by (see Fig. 1)

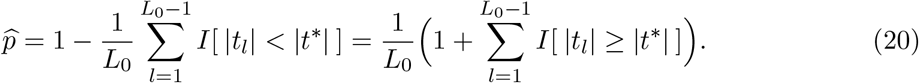

Note that the only possible values for 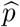 are 1/*L*_0_, 2/*L*_0_,…, 1, which implies that 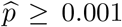 when *L*_0_ = 1000. Moreover, if the significance level of the test is taken to be *a* = *k*/*L*_0_ for some integer *k* such that 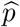 can take value *k*/*L*_0_, then it can be shown that Pr[Type I error] = *a*, whereas if the significance level is taken to be *a* = *k*/*L*_0_ for some integer *k* such that 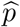 cannot take a value *k*/*L*_0_, then Pr[Type I error] < *a*. Consequently, this procedure can also control the Type I error.

#### Matched pairs group comparison

Assuming availability of *M* pairs of matched values of a given statistical summary from the first and second groups, associated with an analysis region, CpelTdm.jl performs hypothesis testing using the previous randomization testing method, provided that the total number *L* = 2^*M*^ of possible matched group assignments is less than *L*_0_. In this case, the *M* matched pairs are used to form *L* distinct permutations, each containing all *M* pairs but with some group labels being reversed, and a value *t_l_* is computed for the *l*-th permutation using each of the following two differential test statistics:

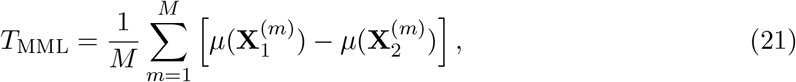

and

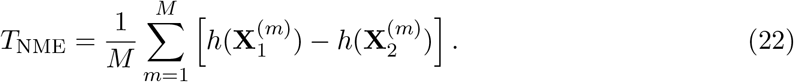

However, if *L* ≥ *L*_0_, CpelTdm.jl performs hypothesis testing using the Monte Carlo based permutation procedure employed in the unmatched case in which the *L*_0_ – 1 values of the test statistic required by the method are determined by independently sampling the set of all *L* matched group permutations with equal probability and by computing the test statistic value for each permutation.

Unfortunately, the previous procedure cannot be used to perform hypothesis testing using the CMD because of its symmetry; i.e., due to the fact that 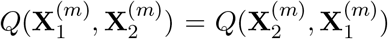. For this reason, CpelTdm.jl simply calculates the average of the CMDs associated with all matched analysis regions, given by 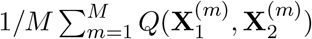, and outputs the results.

Note that, in both the unmatched and matched cases, CpelTdm.jl performs hypothesis testing only when the number of group permutations *L* is large enough (*L* ≥ 20) to produce *P*-values that are not above a 5% significance level (recall that, when *L* < 1000, the *P*-values cannot be less than 1/*L*). Finally, CpelTdm.jl performs multiple hypothesis testing using Benjamini-Hochberg (BH) correction for false discovery rate (FDR) control (see Fig. 1) and provides the adjusted *p*-values in the output.

## References

Abante, J. et al. (2020) Detection of haplotype-dependent allele-specific DNA methylation in WGBS data, Nat. Commun., 11, 5238.

Robinson, M. D. et al. (2014) Statistical methods for detecting differentially methylated loci and regions, Front. Genet., 5, 324.

Hebestreit, K. et al. (2013) Detection of significantly differentially methylated regions in targeted bisulfite sequencing data, Bioinformatics, 29, 1647–1653.

Landan, G. et al. (2012) Epigenetic polymorphism and the stochastic formation of differentially methylated regions in normal and cancerous tissues, Nat. Genet., 44, 1207–1214.

Li, S. et al. (2014) Dynamic evolution of clonal epialleles revealed by methclone, Genome Biol., 15, 472.

Jenkinson, G. et al. (2017) Potential energy landscapes identify the information-theoretic nature of the epigenome, Nat. Genet., 49, 719–729.

Jenkinson, G. et al. (2018) An information-theoretic approach to the modeling and analysis of whole-genome bisulfite sequencong data, BMC Bioinf., 19, 87.

Nielsen, F. (2019) On the Jensen-Shannon symmetrization of distances relying on abstract means, Entropy, 21, 485.

Ernst, M. D. (2004) Permutation methods: A basis for exact inference, Stat. Sci., 19, 676–685.

